# Two-hit mouse model of heart failure with preserved ejection fraction combining diet-induced obesity and renin-mediated hypertension

**DOI:** 10.1101/2024.06.06.597821

**Authors:** Justin H. Berger, Yuji Shi, Timothy R. Matsuura, Kirill Batmanov, Xian Chen, Kelly Tam, Mackenzie Marshall, Richard Kue, Jiten Patel, Renee Taing, Russell Callaway, Joanna Griffin, Attila Kovacs, Dinesh Hirenallur Shanthappa, Russell Miller, Bei B. Zhang, Rachel J. Roth Flach, Daniel P. Kelly

## Abstract

Heart failure with preserved ejection fraction (HFpEF) is increasingly common but its pathogenesis is poorly understood. The ability to assess genetic and pharmacologic interventions is hampered by the lack of robust preclinical mouse models of HFpEF. We have developed a novel “2-hit” model, which combines obesity and insulin resistance with chronic pressure overload to recapitulate clinical features of HFpEF. C57BL6/NJ mice fed a high fat diet for >10 weeks were administered an AAV8-driven vector resulting in constitutive overexpression of mouse *Renin1d*. Control mice, HFD only, Renin only and HFD-Renin (aka “HFpEF”) littermates underwent a battery of cardiac and extracardiac phenotyping. HFD-Renin mice demonstrated obesity and insulin resistance, a 2-3-fold increase in circulating renin levels that resulted in 30-40% increase in left ventricular hypertrophy, preserved systolic function, and diastolic dysfunction indicated by altered E/e’, IVRT, and strain measurements; increased left atrial mass; elevated natriuretic peptides; and exercise intolerance. Transcriptomic and metabolomic profiling of HFD-Renin myocardium demonstrated upregulation of pro-fibrotic pathways and downregulation of metabolic pathways, in particular branched chain amino acid catabolism, similar to findings in human HFpEF. Treatment of these mice with the sodium-glucose cotransporter 2 inhibitor empagliflozin, an effective but incompletely understood HFpEF therapy, improved exercise tolerance, left heart enlargement, and insulin homeostasis. The HFD-Renin mouse model recapitulates key features of human HFpEF and will enable studies dissecting the contribution of individual pathogenic drivers to this complex syndrome. Addition of HFD-Renin mice to the preclinical HFpEF model platform allows for orthogonal studies to increase validity in assessment of interventions.

**NEW & NOTEWORTHY:** Heart failure with preserved ejection fraction (HFpEF) is a complex disease to study due to limited preclinical models. We rigorously characterize a new two-hit HFpEF mouse model, which allows for dissecting individual contributions and synergy of major pathogenic drivers, hypertension and diet-induced obesity. The results are consistent and reproducible in two independent laboratories. This high-fidelity pre-clinical model increases the available, orthogonal models needed to improve our understanding of the causes and assessment treatments for HFpEF.

## INTRODUCTION

Heart failure with preserved ejection fraction (HFpEF) is a complex disease, and despite many advances in research, its underlying mechanisms are not yet fully understood (1). This is due, in part, to the lack of pre-clinical small animal models which faithfully recapitulate the human condition. There is no consensus on the required elements to define rodent HFpEF (2–5). Consequently, current rodent models have notable downsides, including significant lead time hindering rapid experimentation (6), need for surgical expertise (7), predominant hypertensive phenotype without metabolic dysregulation (8), or inability to use genetic manipulation, as in the ZSF-1 rat or leptin signaling mutant models (9–11). Additionally, the best characterized mouse model to date using L-NAME and high fat diet (HFD) (12), induces endothelial dysfunction using a pharmacological inhibitor of vascular NO signaling and some have shown that this model is not consistently reproducible (13, 14).

We sought to devise a complementary “two-hit” model of mouse HFpEF that combines the most common human drivers of the metabolic phenogroup of human HFpEF, diet-induced obesity and chronic hypertension. Among HFpEF patients, the phenogroups with obesity/diabetes, abnormal systemic metabolism, liver and renal dysfunction, and high renin level exhibit the worst overall prognosis, and lowest survival probability (15, 16). Our goal was to establish a mouse model that exhibits both cardiac and extra-cardiac manifestations of HFpEF and meets the following criteria: 1) left ventricular (LV) hypertrophy (LVH) driven by a pathogenic driver relevant to the human condition, 2) obesity and insulin resistance, 3) LV diastolic dysfunction, 4) preserved LV systolic function, and 5) exercise intolerance. We also sought to develop a model with relatively balanced contributions of increased peripheral vascular resistance and obesity with insulin resistance. The model described here, termed “HFD-Renin” mice, involves modest, chronic hypertension achieved by adeno-associated virus overexpression of renin (AAV8-Renin) to activate the renin-angiotensin-aldosterone system (RAAS) in the setting of diet induced obesity (DIO) producing a modest but significant LVH, LV diastolic dysfunction with stable systolic function, exercise intolerance and obesity-related glucose intolerance and insulin resistance. This model utilizes common pathogenic drivers resulting in features of human HFpEF and allows for assessment of the relative contribution of each driver independently (16, 17). Given concerns of reproducibility with prior published models, we developed and conducted confirmatory studies at two sites in a coordinated manner to verify reproducibility. The availability of multiple verified HFpEF models will be valuable to the field, allowing for more rigorous assessment of therapeutic interventions via orthogonal studies.

## MATERIALS AND METHODS

### In vivo studies

Mouse studies were performed in parallel at two separate institutions. Penn utilized male and female C57BL/6NJ mice (Jackson Laboratory, Stock 005304) age 8-10 weeks old, and randomized to chow or 60%kcal HFD (D12492i, Research Diets) arms. Pfizer utilized 20-week-old C57BL/6N male mice fed chow or preconditioned at age 8 weeks for 12 weeks on 60% HFD-fed (Taconic, DIO-B6NTac). Mice were housed in a facility under a 12-hour light/12-hour dark cycles. For tissue collection, mice were fasted for 4 hours prior to anesthesia with intraperitoneal pentobarbital (Sagent, 100 mg/kg). Tissue weights were normalized to tibia length. Lungs and livers were excised and weighed “wet” and, for lungs only, following 48 hours at 37°C “dry”. For empagliflozin (empa) studies, mice were randomized to treatment with drug (Advanced ChemBlocks Inc, Cat#: G-7261) admixed at 500mg/kg (Pfizer) or 50mg/kg (Penn) in 60%kcal HFD, validated to match the human pharmacologic effects (18).

### AAV8-Renin Generation and Treatment

Mouse *Ren1d* cDNA was purchased from Horizon Discovery (Clone ID 30313659, Sequence BC061053) and mutated at F61R and P65S as previously described (to enable prorenin cleavage to renin in non-renal locations, see Supplemental Fig. S1A) using the In-Fusion HD system (Clontech, #639650) (19). Construct was subcloned in the pAAV.TBG.PI.Null.bGH (Addgene #105536) plasmid provided by the Vector Core at University of Pennsylvania. All cloning and mutagenesis primer sets are listed in Supplemental Table S1. pAAV.TBG.PI.mRen1(F61R)(P65S).bGH (AAV8-Renin) or null constructs were packaged at the Vector Core. For the Pfizer model, AAV was produced at the University of Massachusetts Medical Center Gene Therapy Center and Vector Core. 18-20-week-old C57BL6/NJ male mice preconditioned on 10 week of chow or 60% kcal HFD were randomized to receive AAV8-Renin or AAV8-null virus. Mice were anesthetized using 2.5ppm isoflurane prior to injection via the medial canthus into the retro-orbital venous plexus.

Based on extensive dose-response testing, doses were selected to generate a 2-3x increase over basal renin expression but without affecting weight gain (Supplemental Fig. S1). Given viral production occurred in two independent facilities, empiric doses were as follows: Penn lean mice received 3.0×10^10 and HFD-fed mice received 1.0×10^11 viral genomes (vg) diluted in sterile saline, injected in a volume of 100ul. Pfizer lean mice received 1.0×10^10 and HFD-fed 3×10^11 vg. Higher doses of AAV-Renin were required to raise plasma renin levels in obese mice, potentially due to lower transduction efficacy of AAVs in the fatty liver of the HFD model. Blood samples were obtained 1-week post-injection and assayed for circulating renin using the mRen1 DuoSet ELISA (Bio-techne, #DY4277) per manufacturer protocol. Samples from mice receiving AAV8-Renin were diluted 1:50.

### Telemetry study

Radiotelemetry recording of arterial pressure was performed in 8–10-week-old male C57BL/6N mice, implanted with telemeters (PA-C10, Data Science) by Envigo (Invito, Indianapolis, IN). After 1 week, telemetry units were activated, and arterial pressure was recorded for baseline recording. Mice were randomized and dosed with either 1×10^10 AAV null or 3×10^9, 1×10^10 or 3×10^10 AAV-Renin. Two weeks post-AAV dosing, mice were treated with enalapril (200mg/L; Sigma) in the drinking water for 3 days, followed by switching back to normal drinking water.

### Mouse echocardiography

Ultrasound examination of the left ventricle was performed independently at both study sites using a Fujifilm VisualSonics Ultrasound System. At the Rodent Cardiovascular Phenotyping Core of the University of Pennsylvania Cardiovascular Institute, adult mice were anesthetized with an intraperitoneal injection (0.05 mg/g) of 2% avertin (to maintain a heart rate close to 600 beats per minute or higher for the evaluation of LV systolic function). Hair was removed from the anterior chest using chemical hair remover, and the animals were placed on a warming pad in a left lateral decubitus position to maintain normothermia. Ultrasound gel was applied to the chest. Care was taken to maintain adequate contact while avoiding excessive pressure on the chest. 2D long-axis and short-axis M-mode images were obtained. Diastolic function–related parameters were evaluated using a modified 4-chamber view. Transmitral inflow velocities, tissue Doppler, pulmonary vein Doppler and left atrial strain measurements, as previously described (8), were obtained following intraperitoneal injection of zatebradine (funny current inhibitor, 0.008 mg/gm body weight) to reduce HR to 425–475 bpm (15%–25 % HR reduction compared with baseline) when needed. After completion of the imaging studies, mice were allowed to recover from anesthesia and returned to their cages. At Pfizer, animals were anesthetized at 2% isoflurane (chow) or 3% isoflurane (HFD) with heart rate goals 300-450bpm for diastolic parameters and >400bpm for functional parameters. Images were obtained similarly. Imaging and image analysis was performed by blinded sonographers and analyzed using Vevo Lab software (VisualSonics).

### Exercise testing

Mice were exercised to exhaustion utilizing similar protocols. At Penn, mice underwent 3 consecutive days of exercise on a 5-lane treadmill with shock grid (Maze Engineering) starting with 2 days of an acclimation protocol of 9 minutes at 10m/min followed by 1 minute at 20m/min at a grade of 6-8 degrees. Testing was performed on a 25-degree treadmill, starting at 10 m/min for 3 minutes followed by a ramp over 3 minutes to a final speed of 18 m/min, continuing to exhaustion. At Pfizer, animals were acclimated to the Exer3/6 Animal Treadmill (Columbus Instruments) for 2 days prior to testing– Day 1: treadmill belt off and shock grid on for 5min before an additional 5min at 2.5m/min; Day 2: 5min at 2.5m/min followed by another 5min at 5m/min. Testing was performed at a 5 degree incline starting at 5m/min for 2 minutes, increased to 8m/min for 2 minutes and then increased an additional 2m/min for every 2 minutes until the animal was exhausted. Exhaustion was defined by more than 5 seconds in contact with shock pad, or more than 50% of the time off the treadmill.

### Metabolic measurements

Glucose and insulin tolerance testing (GTT and ITT, respectively) was conducted after a 5-hour fast. A bolus of glucose (1 g/kg) or insulin (0.75U/kg) was administered via intraperitoneal injection. A blood sample was obtained from the tail tip for the measurement of baseline glucose using a handheld glucometer (Accu-Chek Aviva Plus, Roche or AlphaTrak2 #71681-01) at 0, 15, 30, 45, 60, and 90 minutes following the injection. At Penn, plasma insulin was measured using Mouse Insulin ELISA kit (Mercodia). Cholesterol and triglycerides were measured using Infinity colorimetric assays (ThermoScientific). Plasma samples at Pfizer were analyzed using Insulin Rodent (Mouse/Rat) Chemiluminescence ELISA kit (ALPCO, #80-INSMR-CH01). HOMA-IR was calculated using the insulin and glucose measurements, using the formula: insulin (µU/ml)*glucose (mmol/L)/22.5. Targeted metabolomics was performed at the Metabolomics Core in the Penn Cardiovascular Institute as previously described, from flash frozen mouse hearts (20).

### Histology

The tissues prepared as previously described (20) including overnight fixation in 4% paraformaldehyde in PBS and dehydration through sequential ethanol washes. The tissues were then embedded in paraffin and sectioned as 6mm. Picro Sirius Red staining was performed by the Histology and Gene Expression Co-Op at the University of Pennsylvania and imaged by the Children’s Hospital of Philadelphia Pathology Core. Collagen staining was quantified with Aperio ImageScope (Leica Biosystems).

### RNA isolation and qRT-PCR

Total RNA was isolated using the RNeasy Mini Kit (QIAGEN) and the RNase-Free DNase Set (QIAGEN) according to the manufacturer’s instructions. cDNA was synthesized using the Affinity Script cDNA Synthesis Kit (Agilent Technologies) with 0.25 μg total RNA. PCR reactions were performed using Brilliant III Ultra-Fast SYBR Green QPCR Master Mix (Agilent Technologies) on a QuantStudio 6 Flex Real-Time PCR System (Applied Biosystems) with specific primers for each gene. The Penn primer sets are listed in Supplemental Table S1. The expression of target mRNAs was normalized by that of *Rplp0* (36B4). Pfizer cardiac gene expression was measured by RT-PCR with TaqMan probes (*Myh6*: Mm00440359_m1, *Myh7*: Mm00600555_m1) normalized to *Hprt* (Mm03024075_m1). All experiments were repeated independently, with samples in either triplicate or quadruplicate.

### RNA-Seq library preparation, sequencing and analysis

RNA library preparations and sequencing reactions were conducted at GENEWIZ, LLC. (South Plainfield, NJ), per their standard operating procedure and as previously described (21, 22). We used Salmon v1.10.2 to count reads in GRCm38 (mm10) reference transcripts (23). The counts were summarized at gene level and differentially expressed genes were identified with DESeq2 1.40.2 (24). Genes with an FDR below 0.05, when comparing conditions of interest, and at least a 1.2-fold expression change (up or down) were considered differentially expressed. We performed pathway enrichment analysis, using gProfiler web service with FDR < 0.05 cutoff and enrichment at least 2 (25).

### Statistics

For 2-group comparisons, a Student’s *t* test was performed when the data were normally distributed based on a Shapiro-Wilk test (α = 0.05). For multiple comparisons, 2-way ANOVA with Tukey’s post hoc test was performed. GraphPad Prism 7.04 or 8.03 (GraphPad Software) was used for graphing and statistical analysis.

### Study approval

All animal experiments were performed in accordance with NIH guidelines for the humane treatment of animals and approved by the Institutional Animal Care and Use Committees of the Perelman School of Medicine at the University of Pennsylvania and Pfizer.

## RESULTS

### Development and optimization of AAV-mediated renin delivery to achieve hypertension-related LV hypertrophy and remodeling in mice

We sought to use an AAV8 delivery system to achieve a mild degree of hypertension in mice. An AAV-Renin vector (details of the construct provided in Methods) or null backbone vector (control) were administered in order to constitutively express a cleavable form of renin in the liver (19). Dose-ranging studies demonstrated a dose-dependent increase in circulating mouse renin concentration (Supplemental Fig. S1A). No change in body weight was observed during the 6-week study period (Supplemental Fig. S1B). Implantable telemetry testing revealed a dose-dependent, modest (20-40mmHg) increase in systolic blood pressure (Supplemental Fig. S1C), which was amenable to treatment with standard of care angiotensin converting enzyme (ACE) inhibitor enalapril. Consistent with the hypertensive state, increased normalized biventricular heart weight, left atrial enlargement, and modest lung congestion developed in the AAV-renin-treated mice (Supplemental Fig. S1D). Gene markers of LV remodeling were also induced including a decrease in the myosin heavy chain 6 (*Myh6*)/*Myh7* ratio (Supplemental Fig. S1E).

The dose ranging studies were repeated in HFD-treated mice. Mice were maintained on HFD for 10-12 weeks starting at 8 weeks of age prior to AAV administration. Dose-dependent LVH was achieved in the context but required higher viral doses presumably due to presumed decreased viral transduction in a fatty liver (Supplemental Fig. S1F). Notably, at the highest doses, significant loss of body weight was observed (Supplemental Fig. S1G). The dose-ranging studies allowed for the selection of similar, though not identical, doses at both institutions (lower for chow-fed animals, higher for HFD-fed animals) that resulted in the optimal 2-3x increase in circulating renin and LVH with minimal perturbance in body weight.

### The HFD-Renin model results in features of HFpEF including systemic metabolic derangements, LV hypertrophy, diastolic dysfunction, and exercise intolerance

Having established that AAV-Renin produces modest hypertension and cardiac hypertrophic remodeling, this effect was overlayed on HFD-fed animals. For the HFpEF model, 8–10-week-old mice pre-conditioned on chow (control) or HFD for 10 weeks were injected with AAV-Renin or null vector (control) (Fig. 1A). Renin dosing was calibrated to achieve a 2-3x increased in circulating protein levels (Fig. 1B). HFD-feeding reliably resulted in obesity, though AAV-Renin dosing at week 10 caused a modest flattening in weight gain (Fig. 1C). The obese mice had impaired glucose tolerance and insulin resistance (Fig. 1D and S2A). This corresponded with fasting hyperglycemia, hyperinsulinemia and hypercholesterolemia (Supplemental Fig. S2B). There were no observed metabolic effects due to AAV-Renin (see chow+Renin cohort).

**Figure 1.**
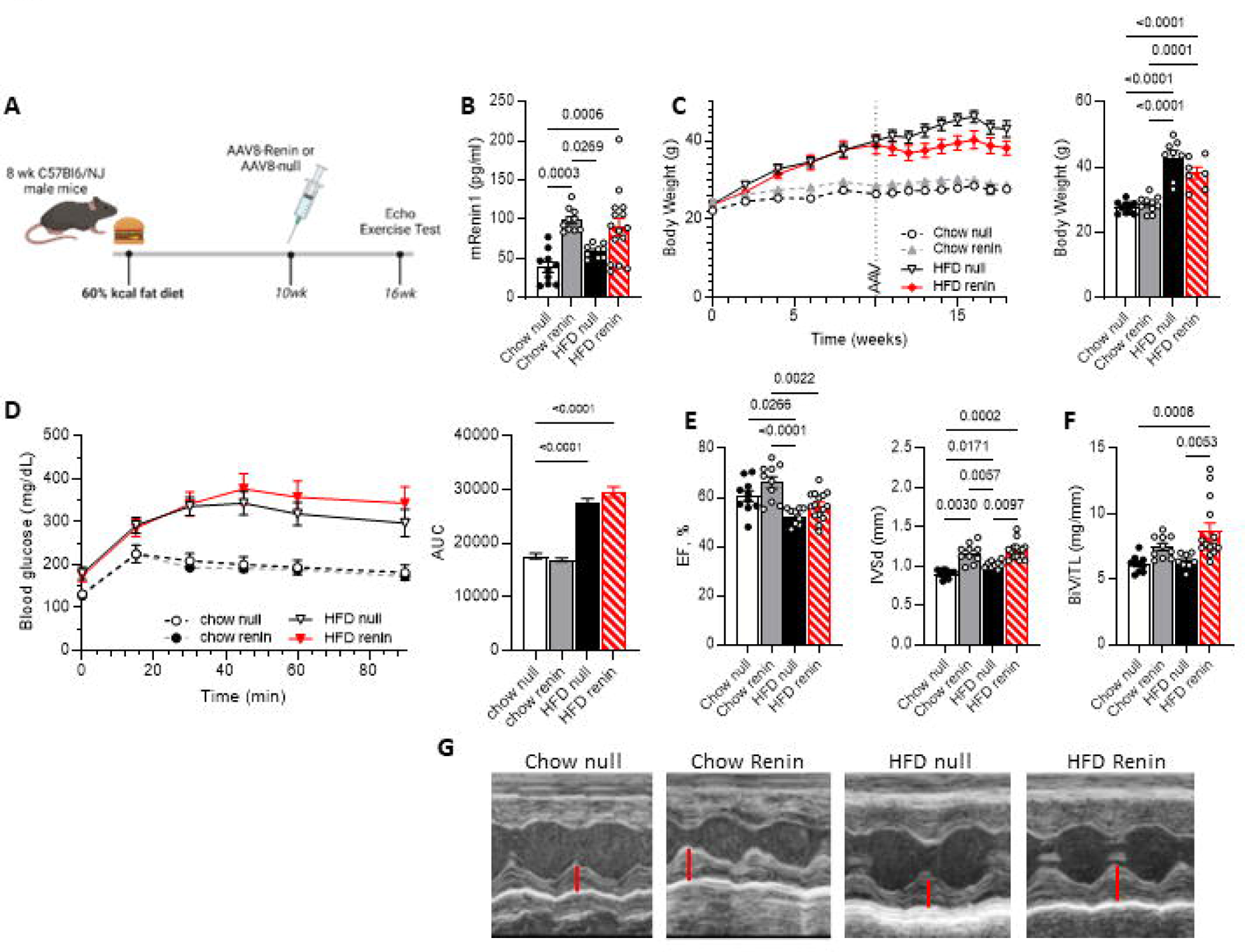
Novel “2-hit” HFpEF mouse model driven by renin overexpression and HFD. A.) Schema of HFD-Renin model, where C57BL/6NJ mice are preconditioned on HFD for 10 weeks prior to injection of AAV8-mRenin with endpoints at week 16-18. Created with Biorender.com. B) Serum renin levels 1-week post-injection. C.) Weight trend (left) and terminal weights (right). D.) Glucose tolerance test with associated area-under-curve (right) in 4-hour fasted male littermate mice. E.) Echocardiographic measures at 16 weeks: LV ejection fraction (EF) and interventricular septal thickness in diastole (IVSd). F.) Tibia length-normalized gross biventricular weight (BiV/TL). G.) Representative m-mode echocardiograms, red bar notes LV posterior wall dimension in diastole. N= 5-9 per conditions. Data displayed as mean ± SEM. Statistical analyses used one-way analysis of variance (ANOVA) followed by multiple-comparisons test. P values < 0.05 displayed on graphs.

All cohorts had stable LV systolic function at 6 weeks post-injection (Fig 1D and Supplemental Table S2). AAV-Renin resulted in 30-40% increase in LV intraventricular septal wall thickness (IVSd) and LV mass index (LVMI) as determined by echocardiography (Fig. 1D-F, and Supplemental Table S2). Normalized ventricular weights (biventricular weight to tibia length, BiV/TL) were similarly increased, with an additive effect between HFD and renin expression (Fig. 1E). Despite the LVH, there was no significant change in ventricular volumes (Supplemental Table S2).

To assess diastolic function, multiple complimentary endpoints were utilized. HFD or AAV-Renin alone did not produce significant changes in echo-based parameters. An additive effective of HFD and AAV-Renin was observed in mitral valve inflow and tissue doppler imaging parameters (increased E/e’, a trend in Tei index, no change in E/A ratios, Fig. 2A and 2B) as well as global longitudinal and radial strain and strain rates (Fig. 2C and Supplemental Table S2). Radial strain appeared to be a more sensitive marker of dysfunction in this model. As observed in the human HFpEF condition, the mouse left atria (LA) was impacted including significantly increased in normalized LA weight relative to the control (Fig. 2D) and trended towards increased LA volume measured by echo (Fig. 2E), consistent with LV diastolic dysfunction. LA strain, a standard human clinical parameter, was reduced. Similarly, pulmonary vein doppler flow was significantly decreased in the HFD Renin cohort, indicative of decreased LV compliance (Fig. 2F). Expression of elevated natriuretic peptides *Nppb* and *Nppa,* a diagnostic criterion in human disease, were significantly elevated in heart tissue from HFD Renin animals compared to other treatment groups (Fig. 2G). Notably, lung congestion was observed in the Pfizer (Supplemental Fig. S1D) but not in the Penn cohort (Supplemental Table S2).

**Figure 2.**
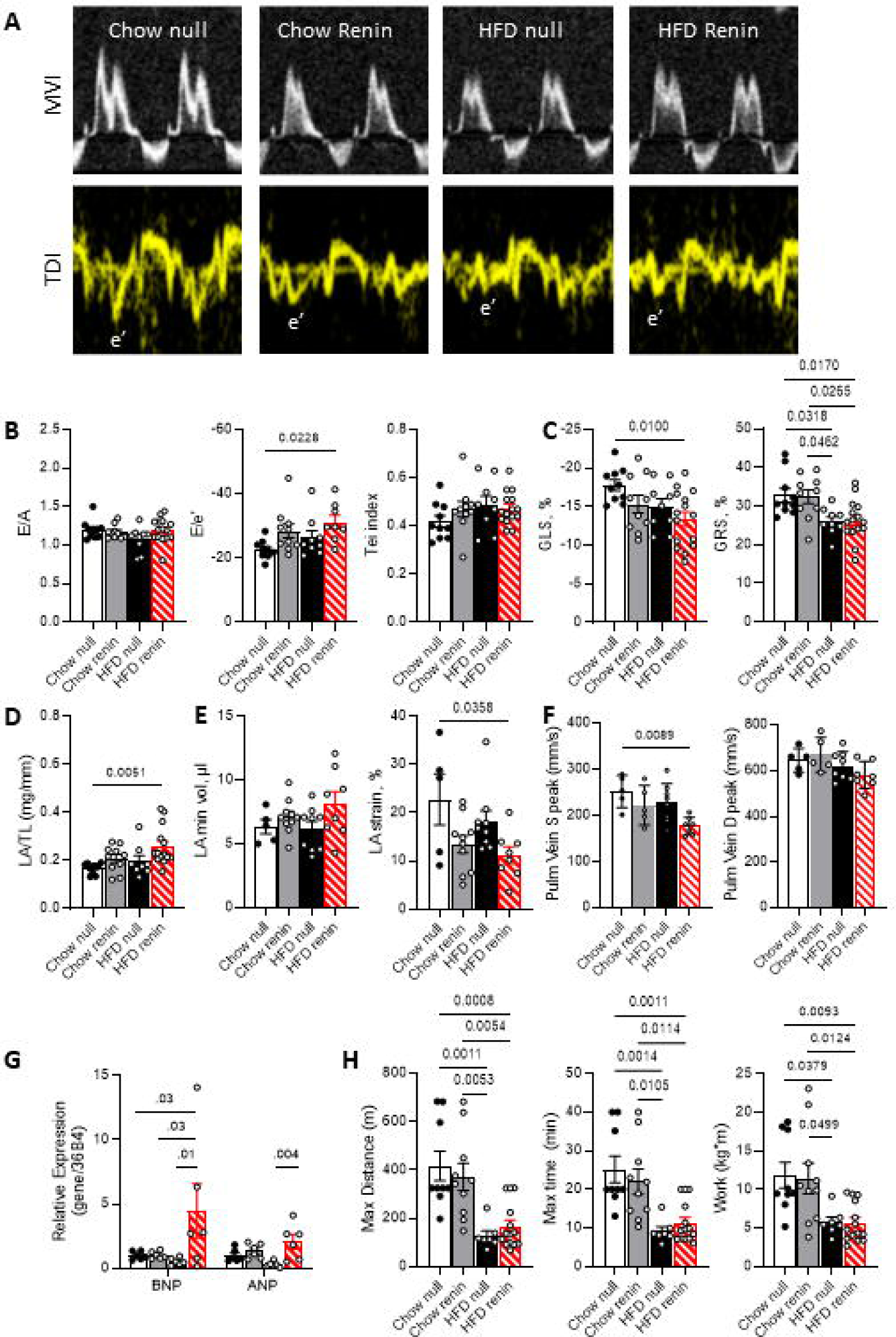
HFD-Renin HFpEF model exhibits LV diastolic dysfunction, abnormal LA function, and exercise intolerance. A.) Representative echocardiographic mitral valve inflow and tissue doppler images with B.) measurements of diastolic function including E/A and E/e’ ratios, and Tei index (n=5-8). C.) Echocardiographic left ventricular global longitudinal and radial strain (n=4-8). D.) Tibia length-normalized left atrial (LA) gross weight. Representative E.) LA minimum volume and reservoir strain, and F.) Doppler echo-measured pulmonary vein flows during systole (S) and diastole (D). G.) QT-PCR from bulk ventricle RNA for indicated genes. H.) Treadmill exercise testing to exhaustion. Data displayed as mean ± SEM. One-way analysis of variance (ANOVA) followed by multiple-comparisons test. P values < 0.05 displayed on graphs.

Exercise intolerance is the most consistent symptom in patients with HFpEF, though the etiology is debated (26, 27). Mice underwent exercise testing to exhaustion to determine the combinatorial effects of DIO and hypertension. HFD Renin animals displayed greatly diminished exercise tolerance in both time and distance to exhaustion as well as total work, which adjusts for body mass differences (Fig. 2G). Chow AAV-Renin mice did not perform differently from control mice. Accordingly, the dominant effect was due to HFD, as has been demonstrated in prior studies (7).

HFpEF is more prevalent in human females, yet as a group, are relatively understudied in clinical trials (28). We tested the cumulative effects of HFD and AAV-Renin in a cohort of female mice using the same protocol as that for males (Supplemental Fig. S3A and S3B). Female mice fed HFD became obese, but to a lesser extent than male mice, consistent with the pre-clinical rodent literature (29–31). The HFD-fed groups developed insulin resistance (data not shown). Renin-treated groups had preserved systolic function (Supplemental Fig. S3C). Both HFD and AAV-Renin resulted in mild LVH with an additive degree (20-30%) in the HFD-Renin cohort (Supplemental Fig. S3D). Diastolic parameters were mixed: echo parameters for diastolic dysfunction, including E/A, E/e’, and IVRT, were unchanged in the HFD-Renin group compared to controls (Supplemental Fig. S3E). However, decreased pulmonary vein doppler flow and LA enlargement, consistent with decreased ventricular compliance, were observed in the HFD Renin group (Supplemental Fig. S3F and S3G). Lastly, female HFD-Renin mice had decreased exercise performance in an additive manner (Supplemental Fig. S3H). Taken together the HFpEF phenotype driven by HFD-Renin was milder in female mice.

### Two-hit HFpEF model results in cardiac fibrosis

Cardiac fibrosis is a hallmark of human HFpEF, and variably demonstrated in current models (6, 8, 12). Analysis of histologic sections of LV myocardium free wall myocardium stained with picrosirius red demonstrated a trend towards increased collagen deposition in the HFD-Renin group (Fig. 3A-B). Perivascular changes were more apparent than interstitial fibrosis, similar to the known effects of angiotensin II in rodents (8). These observations correlated with increased expression of markers associated with fibrosis, fibroblast activation, and extracellular matrix production by QT-PCR (Fig. 3C).

**Figure 3.**
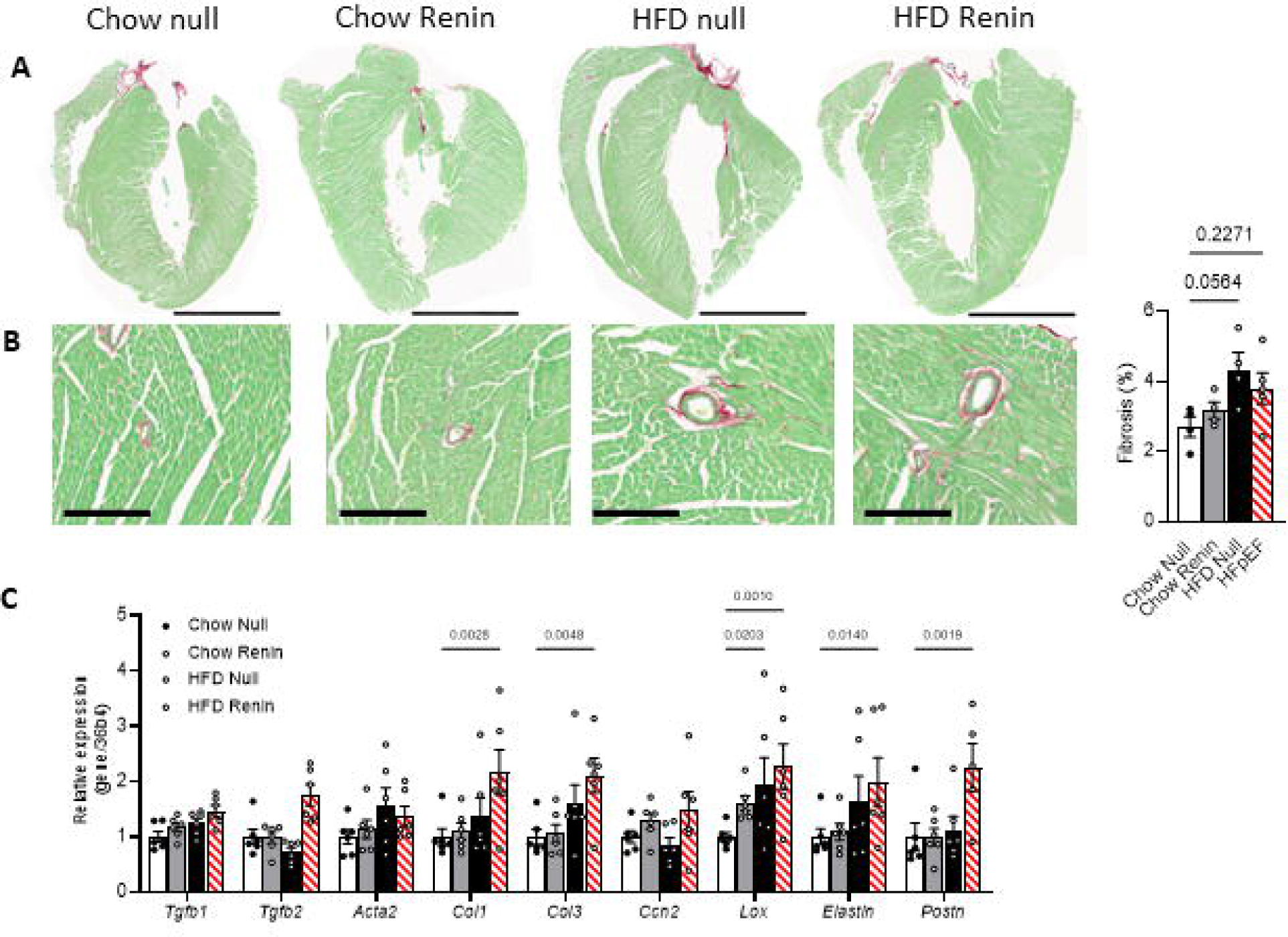
HFD-Renin HFpEF results in increased cardiac fibrosis. Representative histology with picrosirius red and fast green counterstain of A.) whole heart (scale bar 3mm) and B.) 20x magnification (scale bar 200µm) demonstrating perivascular fibrosis, quantified from 2 separate regions in n=4 independent samples. C.) QT-PCR from cardiac tissue for gene markers of fibrosis and myofibroblast activation, n=6. Data displayed as mean ± SEM. One-way analysis of variance (ANOVA) followed by multiple-comparisons test. P values < 0.05 displayed on graphs.

### Transcriptomic and metabolomic changes mirror observations in human HFpEF

Targeted gene expression changes suggested cardiac remodeling largely predicated on fibrotic and hypertrophic pathways (Fig. 2H, 3C). To further understand the gene expression signature of the HFD-Renin model, we performed bulk RNA sequencing (RNA-seq) of LV samples from control, DIO, hypertensive and HFpEF samples. Principal component analysis and heat mapping suggested that the null-treated chow and HFD groups were notably similar, whereas the chow Renin group was quite distinct (Fig. 4A and 4B). The addition of HFD to renin caused a left shift, repositioning the HFpEF group to overlay with control and HFD only groups. We analyzed the effects of the individual drivers and combined effect on differentially expressed genes (DEGs). DIO primarily drove downregulation of genes associated with metabolic pathways and cell signaling (839 downregulated DEGs with adjusted p-value < 0.01) (Supplemental Fig. S4A). Conversely, AAV-Renin was responsible for a large upregulated on genes (927 DEGs) involved with proliferative growth and hypertrophic signaling pathways (Supplemental Fig. S4B). Interestingly, several pathways moved in opposing directions (e.g. cAMP signaling, calcium signaling, and circadian entrapment downregulated by HFD and upregulated by Renin). Consequently, some of these pathways were not present in the gene ontology and KEGG pathway analyses in the HFD-Renin condition. Compared to Chow null, the HFD-Renin cohort resulted in a unique set of 498 upregulated and 472 downregulated genes (Fig. 4C), with a separate set of differentially regulated pathways from either the DIO or hypertensive components. This suggests an additive effect, resulting in downregulated metabolic pathways and upregulation of ECM and cardiac hypertrophic growth (Fig. 4D). These signatures were consistent with published transcriptomic data from HFpEF patients (17). Top DEGs are listed in Supplemental Figure S4C.

**Figure 4.**
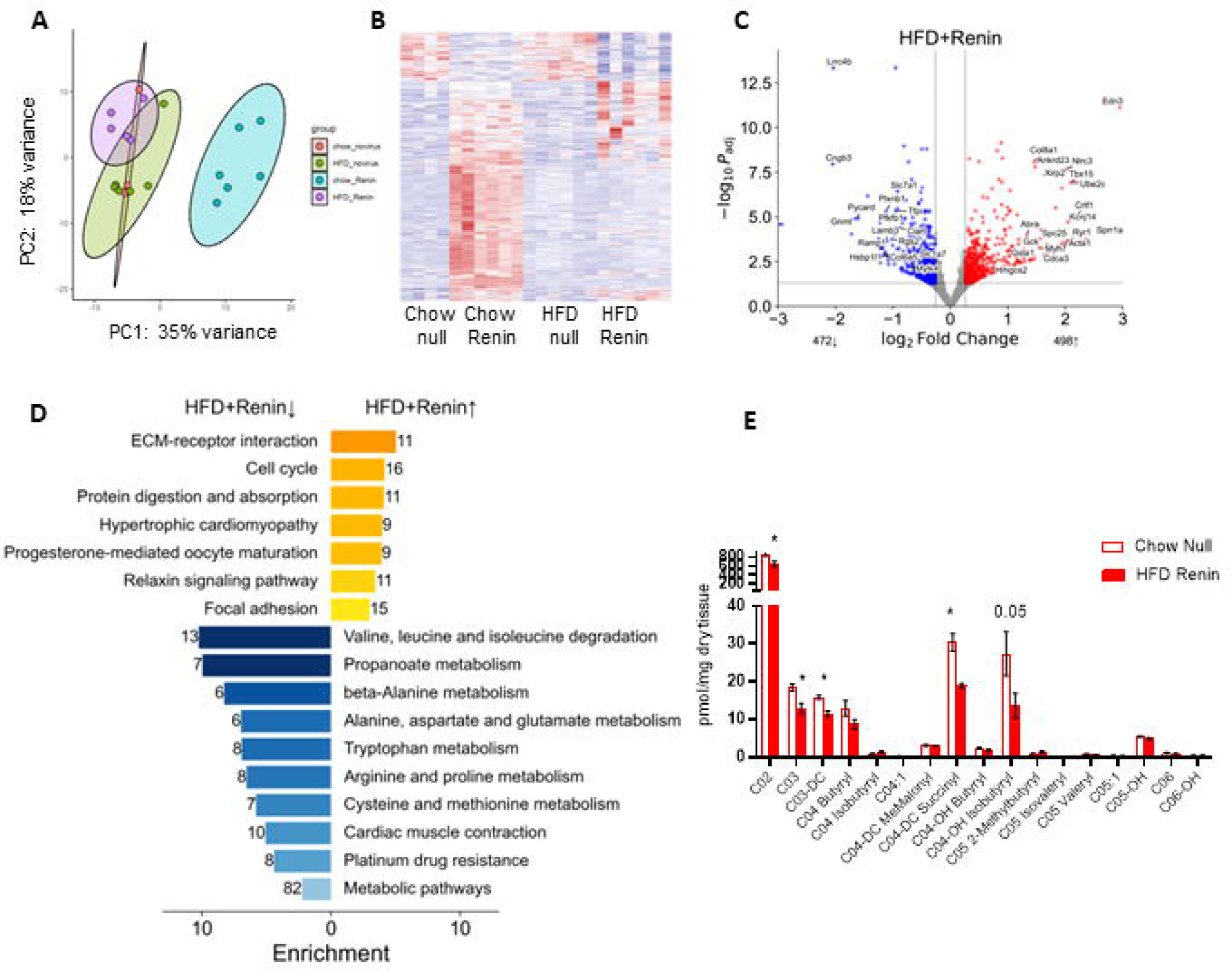
Cardiac transcriptomic and metabolomic profiling of HFpEF hearts shows strong signatures of reduced mitochondrial fuel (BCAA) and activation of ECM/fibrosis. A.) Principal component (PC) analysis and B.) heatmap of ventricular RNAseq comparing control and HFD-Renin groups (n=5-6). C.) Volcano plot of differentially expressed genes in HFpEF (vs control) hearts (n=5-6). D.) KEGG pathway analysis of upregulated (yellow bars) and downregulated (blue bars) pathways. E.) Targeted metabolomics from heart tissue displaying short-chain acylcarnitine species (n=5-7). Data displayed as mean ± SEM. Unpaired, 2-tailed Student’s t-test, with p values as displayed or * <0.05.

Given the observed changes in mitochondrial pathways, including downregulation of branch chain amino acid metabolism and oxidative phosphorylation in the HFD-Renin state, we performed targeted metabolomic analysis in control and HFpEF hearts. Short-chain acylcarnitines, including C03-DC and C04-OH isobutryl, intermediates in BCAA catabolism, were decreased, consistent with the observed reduction in BCAA degradation pathways in the RNA-seq dataset (Fig. 4E). There were few changes in organic acids or medium- and long-chain acylcarnitines (Supplemental Fig. S5).

### Human HFpEF standard-of-care therapy improves a subset of pathophysiologic parameters in the HFD-Renin model

Given that the HFD-Renin model demonstrated changes in metabolism, cardiac remodeling, and exercise tolerance endpoints that capture critical translatable features in HFpEF patients, we next assessed the effects of the newly approved HFpEF therapy sodium glucose cotransporter 2 inhibitor (SGLT2i) empa (32) in this model. Empa was administered concurrent with AAV injection for a total of 14 weeks of treatment at the Pfizer site (Fig. 5A). Treatment with SGLT2i resulted in a slight but significant improvement in body weight, a trend towards reduced blood glucose, with significant reduction in hyperinsulinemia/HOMA-IR (Fig. 5B). No differences were observed in systolic function (Fig. 5C). There was a reduction in echocardiographic LVH and normalized ventricular gross weight following empa therapy (Fig. 5D). Radial strain and echocardiographic measures of diastolic dysfunction were largely unchanged (e.g. IVRT, Fig. 5E), but there was a trend towards diminished normalized LA mass which correlated with reduced liver congestion. Exercise testing demonstrated significant improvement in maximal distance run prior to exhaustion (Fig. 5F). Replication studies conducted in parallel at Penn with a shorter time course of treatment and lower dose demonstrated similar findings (Supplemental Fig. S6). These results suggest that SGLT2i may exert a positive benefit through effects on improved metabolic state, cardiac hypertrophic remodeling, and exercise tolerance and underscore the importance of a model that exhibits multiple pathophysiological responses in order to determine the impact of a given intervention.

**Figure 5.**
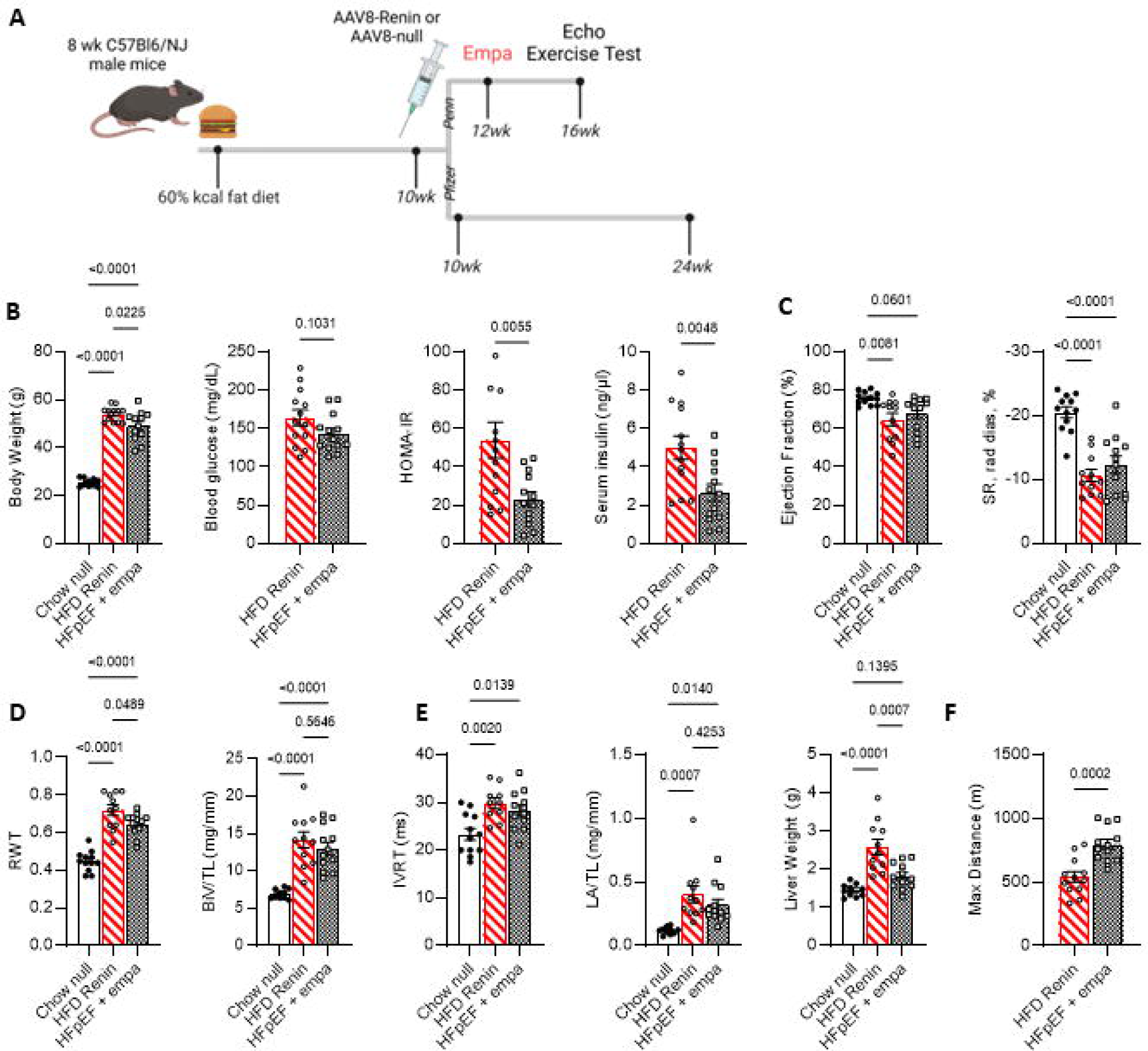
SGLT2 inhibitor treatment improves HFpEF in mice. A.) Schema for empa treatment regimen with HFD-Renin HFpEF model conducted at two independent research sites. B.) Biophysical parameters of Pfizer cohort, n=12-13. C.) Echocardiographic measures of systolic function and strain. D.) LV hypertrophy measures by echo and normalized biventricular weight. E.) Diastolic parameters including IVRT, atrial enlargement and liver congestion. F.) Maximal exercise testing, n=5-11. Data displayed as mean ± SEM. One-way analysis of variance (ANOVA) followed by multiple-comparisons test. P values < 0.05 displayed on graphs. Abbreviations: RWT, regional wall thickness; IVRT, isovolumic relaxation time; SR, strain rate.

## DISCUSSION

Understanding the underlying pathophysiology and identifying new treatment strategies for HFpEF has been hindered, in part, by a lack of mouse models of HFpEF that faithfully recapitulate the human disease state. This challenge is further complicated by the existence of multiple human HFpEF phenogroups necessitating multiple pre-clinical models. The ease of genetic manipulation in mice allows rigorous testing of new targets using genetic loss-of-function approaches complemented by pharmacologic interventions. The two most common drivers in human HFpEF are obesity and HTN—each to varying extent—ranging from obesity with dominant hypertension to modest chronic HTN/renal and vascular dysfunction. We developed a “two-hit” HFpEF model which combines persistent, mild hypertension through renin overexpression with metabolic dysfunction from DIO to simulate the human phenotype. A unique feature of these studies is that the models were assessed in parallel at two separate facilities, increasing confidence in the results and reproducibility for the model. Following careful dose assessment, AAV-Renin could be administered to reliably effect a 20-40mmHg increase in systolic blood pressure and ∼30% LV hypertrophic response without weight loss and preserved systolic function.

We comprehensively phenotyped this new HFpEF model for cardiac and extra-cardiac physiologic endpoints, confirming obesity, insulin resistance, hypertension, preserved systolic function, concentric cardiac hypertrophy, left atrial enlargement, diastolic dysfunction, elevated cardiac natriuretic peptide gene expression, and impaired exercise capacity. To assess diastolic dysfunction, we relied on a plurality of endpoints, including novel left atrial and classic mitral valve echocardiographic measures, gross and normalized tissue weights, exercise intolerance, and natriuretic peptide expression. Multiple metrics of abnormal diastology were present (ventricular strain, E/e’, increased trend in Tei index and IVRT), as well as novel echocardiographic parameters quantifying LA strain, LA volume and pulmonary vein doppler patterns, which have been shown to robustly correlate with diastolic dysfunction (8). Increased hepatic and pulmonary congestion and elevated BNP, a marker of cardiac congestion, mirror the exercise intolerance critical to the human disease. The need for a plurality of markers is not merely pedantic given that E/A and E/e’ are an unreliable sole representation of diastolic dysfunction in mice (33), requiring technical expertise to obtained heart rates slow enough to see E and A separation but not so slow as to affect ventricular function and loading conditions. In addition, multiple pathophysiological endpoints also allow for assessment of therapies that may only impact a subset as we observed with intervention of SGLT inhibitor therapy. We acknowledge the model is limited by modest sex differences, where male mice have greater evidence of HFpEF compared to female mice. This observation is disparate from the human condition but is similar to other mouse HFpEF models (10, 34).

This model allows for assessing how individual pathophysiological interventions that are known drivers (LV pressure overload, caloric excess, abnormal neurohormonal axis) contribute to the development of HFpEF. The ability to isolate component drivers is a powerful technical aspect to the model, querying the specific and additive contributions of hypertension and metabolic dysfunction to the overall HFpEF phenotype. For some metrics, the hypertensive component was the dominant driver, such as measures of LA size and LV hypertrophy. A primary dietary effect was seen in assessing exercise capacity, where HFD largely determined response to exercise. However, for many endpoints, including hypertrophy, diastolic dysfunction, and fibrosis, a significant additive effect was observed beyond either constituent effect.

Ventricular transcriptomic interrogation (RNAseq) was used to assess component effects, identify signature transcriptional changes in the HFpEF state and allow for comparison with recent published human data to further assess relevance (17). Gene expression changes caused by the individual drivers had incomplete similarity to the patterns seen in the HFpEF state, further emphasizing the “syndrome” aspect to this disease – multiple, multiorgan, chronic inputs are required. Among the top upregulated genes in both datasets, many are implicated in extracellular matrix remodeling, cardiac fibrosis and inflammation (e.g. *Acta1, Edn3, Col8a1, Postn, Adamtsl2*). For example, *Acta1* has previously been demonstrated to be upregulated in mouse HFpEF LV (35). Cartilage intermediate layer protein 1 (*CILP1*), reported as a novel sensitive biomarker for cardiac fibrosis (36–38), and disintegrin-like and metalloproteinase domain with thrombospondin type 1 motifs-like (*Adamtsl2*), an extracellular matrix glycoprotein upregulated in human heart failure, act to modulate TGFβ signaling in cardiac fibrosis (39). There was also strong evidence for altered mitochondrial function, including diminished branched chain amino acid catabolism. Targeted metabolomics from cardiac tissue bore out the decrease of BCAA metabolites in the HFpEF state, which is similar to observed changes in human HFpEF samples (40).

To test the translational utility of this HFpEF model, mice were treated with empa, a potent sodium-glucose co-transporter 2 inhibitor (SGLT2i) which has demonstrated improvement in reducing heart failure hospitalization in HFpEF (41, 42). Preclinical data has shown variable benefit of SGLT2i in mouse models of HFpEF, including improvement of fibrosis (43), improved hypertrophy with (44) or without improved diastolic dysfunction (7), and improved electrophysiologic parameters (10). We demonstrate consistent, albeit subtle improvement in only a subset of cardiac and extra-cardiac endpoints, including body weight, insulin sensitivity, LA and LV hypertrophy, and exercise performance, with trends towards improved strain. The benefit of a model that exhibits multiple phenotypic features of HFpEF is emphasized by these results given that therapeutic interventions will likely impact only a subset of the many drivers of this disease. These results highlight the value of this model for translational and mechanistic studies including the potential for genetic interventions that can provide insight into the relevant pathogenic origins of HFpEF.

In summary, we present a tractable, consistent model of mouse HFpEF which recapitulates multiple concerted endpoints observed in the human condition. Compared to the currently available models, this model utilizes AAV-Renin to create a stable, consistent hypertension not affected by food or water intake, or time-limited drug dosing via implantable minipumps (6, 8, 12, 45). Moreover, renin-driven hypertension is a well-known driver of human hypertension and cardiac remodeling. Additionally, there is no imposed vascular dysfunction from an exogenous chemical, such as L-NAME. Hijacking the renin-angiotensin-aldosterone system as a tool to induce hypertension is physiologically relevant to human HFpEF patients, mimicking the significant increase of plasma renin levels observed in the TOPCAT HFpEF clinical trial phenogroup 3 patients (15). Accepting the limitation of using rodents to model human disease, we believe this model, used in conjunction with others previously developed, will allow for critical investigation into a complex and poorly understood form of heart disease.

## DATA AVAILABILITY

All data have been deposited in NCBI’s Gene Expression Omnibus Series accession number GSE269053. The following link has been created to allow review prior to publication: https://www.ncbi.nlm.nih.gov/geo/query/acc.cgi?acc=GSE269053. Please use the following secure token to access the site: wfeveyysthwdxcf.

## SUPPLEMENTAL MATERIAL

Supp. Table 1. Primer sequences.

Supp. Table 2. Echocardiographic and gross morphologic parameters for Renin HFpEF model.

Supp. Fig. 1. Dose titration of AAV8-Renin in lean and obese mice

Supp. Fig. 2. HFD Renin mice developed insulin resistance.

Supp. Fig. 3. Female sex offers protection from protection against HFpEF phenotype in C57Bl/6 mice.

Supp. Fig. 4. Individual drivers of HFpEF have additive transcriptomic changes.

Supp. Fig. 5. Metabolomic profile of HFpEF hearts

Supp. Fig. 6. Effect of SGLT2i on mouse HFpEF (Penn).

## ACKNOWLEDGMENTS

We thank Teresa C. Leone, and Ling Lai at Penn and Federico Damilano at Pfizer for technical assistance. Echocardiography was performed by the Rodent Cardiovascular Phenotyping Core (RRID: SCR_022419) at the University of Pennsylvania supported by the Penn Cardiovascular Institute and NIH S10OD016393. AAV8 virus was generated by the Penn Vector Core (RRID: SCR_022432). Metabolomics studies were performed by the Metabolomics Core (RRID:SCR_022381) in the Penn Cardiovascular Institute and supported in part by NCI P30 CA016520 and NIH P30DK050306. Biorender.com was used for creation of figure schematics.

## AUTHOR CONTRIBUTIONS

JHB, YS, DH-S, RM, BBZ, RRF and DPK conceived and designed research; JHB, YS, KT, XC, MM, RK, TRM, JP, RT, RC and JG performed experiments; JHB, YS, TRM, KB and JG analyzed data; JHB, YS, RRF, and DPK interpreted results of experiments; JHB, KP, RT, and KB prepared figures; JHB, YS, TRM drafted manuscript; and AK, RRF, and DPK edited and revised manuscript. All authors approved the final version of manuscript.

