## Supplementary material for "Two-hit mouse model of heart failure with preserved ejection fraction combining diet-induced obesity and renin-mediated hypertension": Supp Figures

**Supplemental Figure 1. Dose titration of AAV8-Renin in lean and obese mice**. A.) Schema of AAV-B-mRenin1 viral vector. B.) Serum Renin levels 1 week post injection and body weight trends in lean mice (n=3-5) in response to a range of AAV8-Renin doses. C.) Systemic blood pressure measurements and response to enalapril treatment in control and range of AAV8-Renin doses (n=3-4). D.) Tibia length-normalized biventricular weight, left atrial and wet lung weights 6 weeks post injection. E.) QT-PCR markers of cardiac remodeling. F.) Dose-response serum renin levels and normalized biventricular weights and G.) body weight trends in obese mice (n=5-8). Data displayed as mean ± SEM. One-way analysis of variance (ANOVA) followed by multiple-comparisons test. P values < 0.05 displayed on graphs.

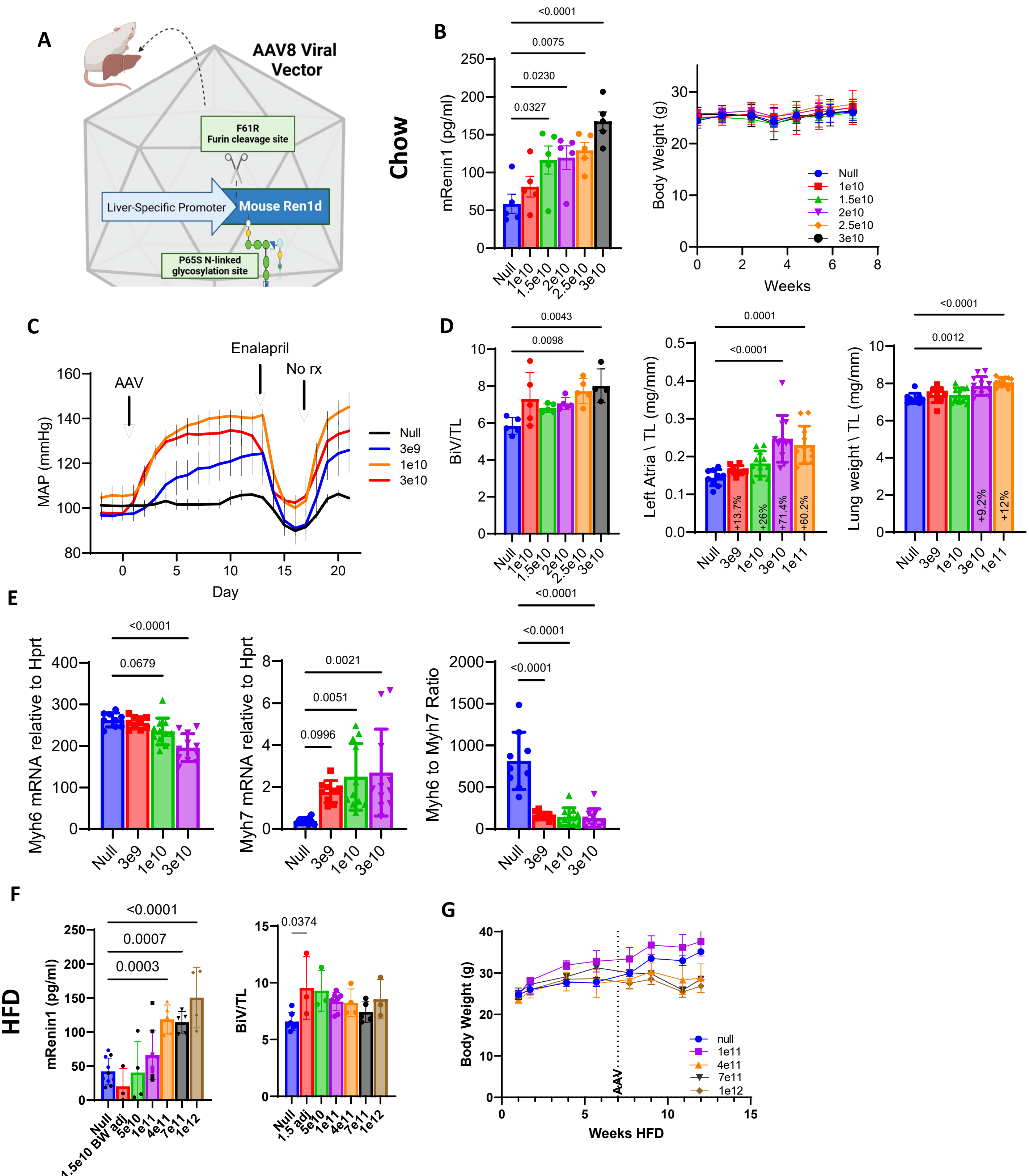

**Supplemental Figure 2. HFD-fed cohorts developed insulin resistance.** A) Insulin tolerance test with associated area-under-curve in 4-hour fasted male littermate mice, n=6-8. B) Fasting serum glucose, insulin, cholesterol and triglycerides, n=5-6 per condition. Data displayed as mean ± SEM. One-way analysis of variance (ANOVA) followed by multiple-comparisons test. P values < 0.05 displayed on graphs.

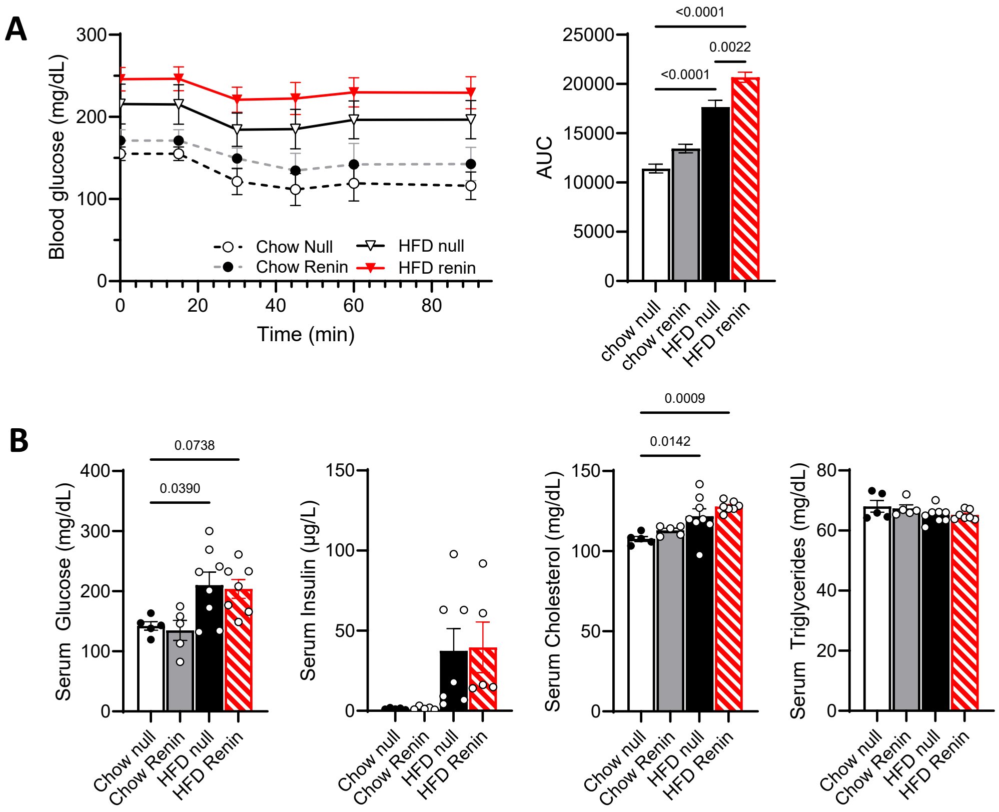

**Supplemental Figure 3. Female sex offers protection against HFpEF phenotype in C57BL/6NJ mice.** A.) Weight trends of female mice undergoing same protocol as in Fig. 1A (n=7-9). B.) Serum renin levels 1-week post-injection. Echocardiographic measures at 16 weeks of C.) systolic function (EF and GRS), D.) LV measurements (intraventricular diastolic septal thickness, IVSd; and biventricular weight normalized to tibia length, BiV/TL), and E.) diastolic function (E/A, E/e', and IVRT, isovolumic relaxation time) . Additional markers of diastolic function include F.) pulmonary vein doppler, G.) gross LA and lung weights, and H.) exercise performance (work). Data displayed as mean ± SEM. One-way analysis of variance (ANOVA) followed by multiple-comparisons test. P values < 0.05 displayed on graphs.

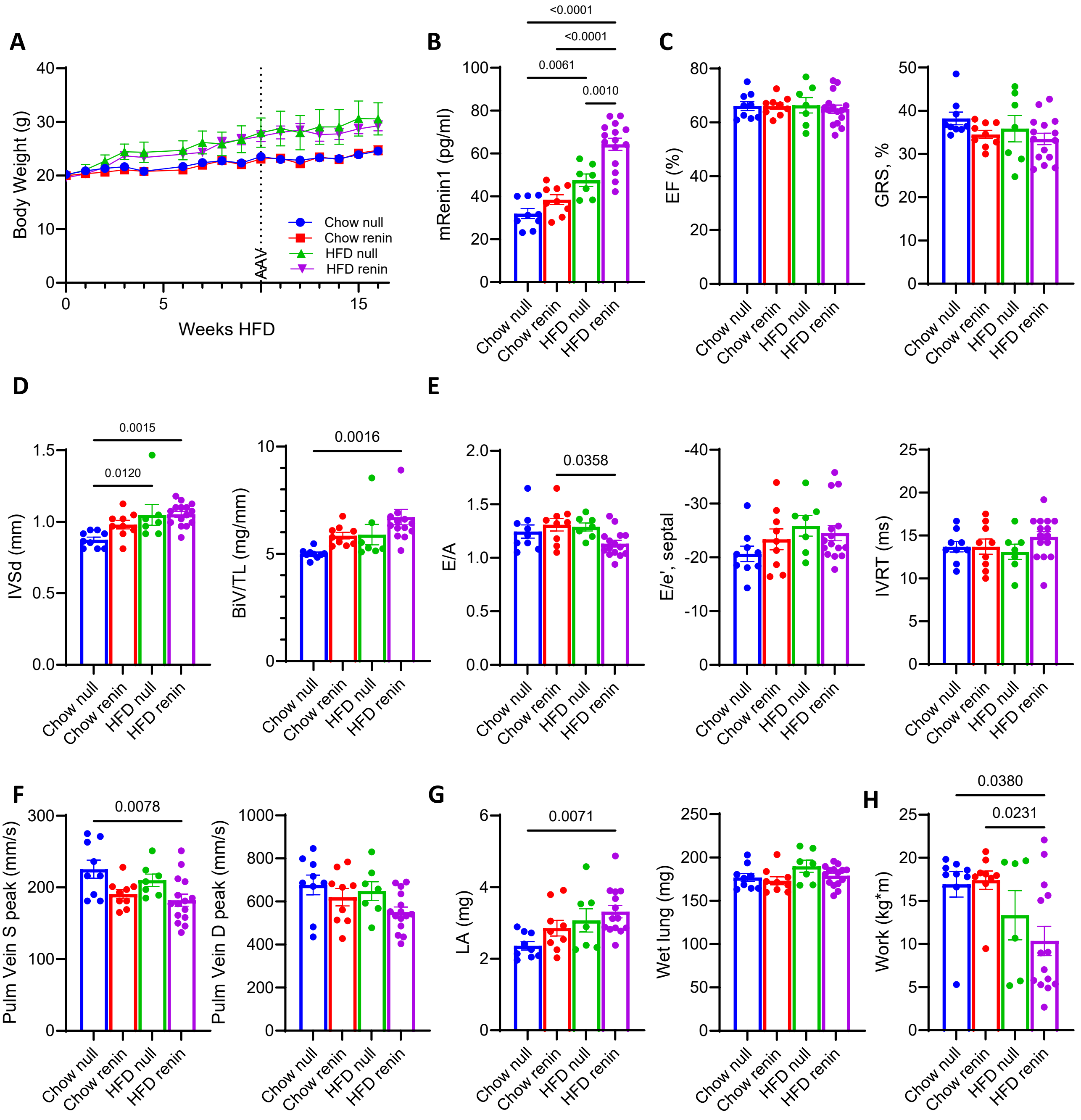

**Supplemental Figure 4. Individual drivers of HFpEF have unique transcriptomic changes.** Volcano plot of differentially expressed genes and selected KEGG analysis showing upregulated (yellow bars) and downregulated (blue bars) pathways due to A.) DIO and B) AAV-Renin (hypertension) effects. N=5-7 C.) Top DEGs by fold-change in HFD+Renin hearts, compared to control.

**A**

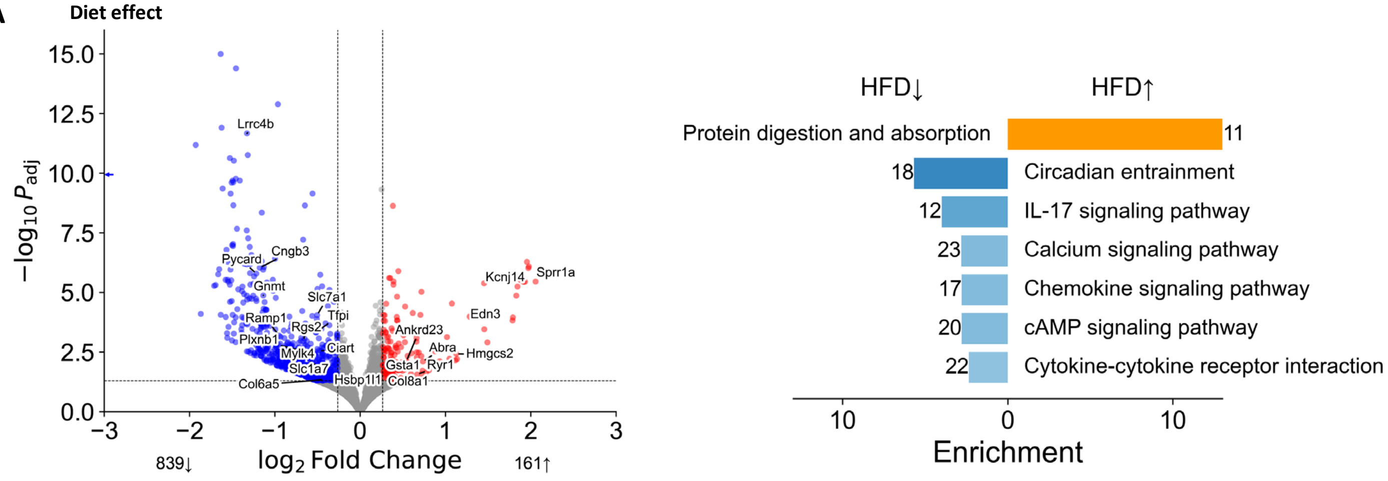


**B**

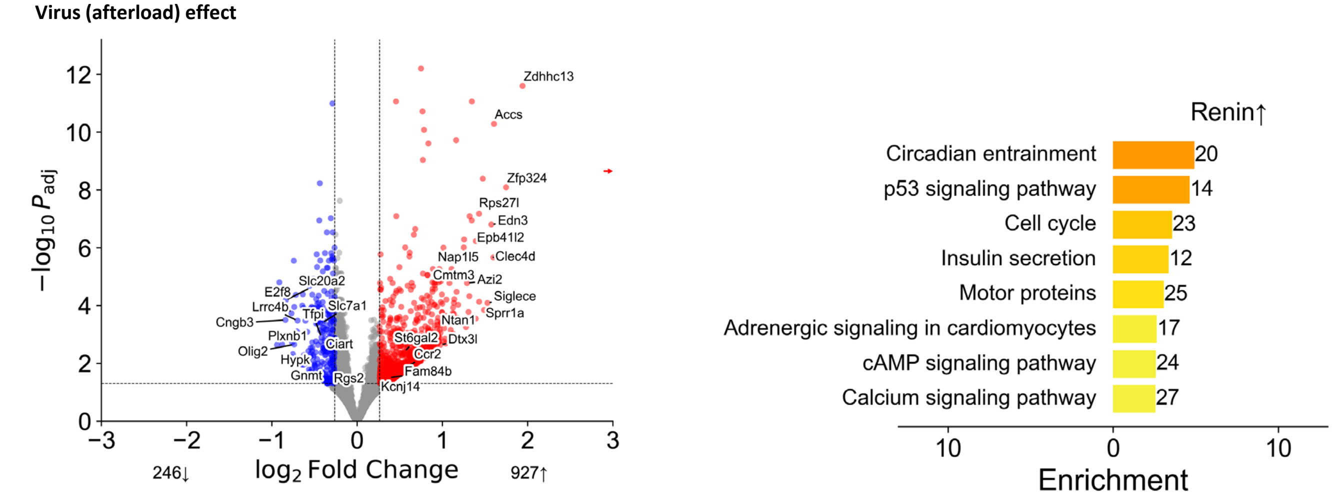


**C**

| Gene ID | Gene name | log2FC | Padj |
|---|---|---|---|
| Sprr1a | small proline-rich protein 1A(Sprr1a) | 4.279211 | 0.000417 |
| Edn3 | endothelin 3(Edn3) | 2.960241 | 6.44E-15 |
| Kcnj14 | potassium inwardly-rectifying channel, subfamily J, member 14(Kcnj14) | 2.031362 | 3.07E-05 |
| Ube2c | ubiquitin-conjugating enzyme E2C(Ube2c) | 1.972274 | 9.14E-06 |
| Crlf1 | cytokine receptor-like factor 1(Crlf1) | 1.83857 | 0.000762 |
| Nlrc3 | NLR family, CARD domain containing 3(Nlrc3) | 1.816272 | 9.28E-06 |
| Cdca3 | cell division cycle associated 3(Cdca3) | 1.799539 | 0.000272 |
| Tbx15 | T-box 15(Tbx15) | 1.769236 | 0.000285 |
| Acta1 | actin alpha 1, skeletal muscle(Acta1) | 1.75458 | 0.001469 |
| Ryr1 | ryanodine receptor 1, skeletal muscle(Ryr1) | 1.727095 | 0.001845 |
| Vgll2 | vestigial like family member 2(Vgll2) | 1.712903 | 0.002431 |
| Shisa3 | shisa family member 3(Shisa3) | 1.652117 | 0.000162 |
| Lman1l | lectin, mannose-binding 1 like(Lman1l) | 1.646589 | 0.002341 |
| Spc25 | SPC25, NDC80 kinetochore complex component, homolog (S. cerevisiae)(Spc25) | 1.522328 | 0.000431 |
| Vil1 | villin 1(Vil1) | 1.480644 | 0.003582 |
| Xirp2 | xin actin-binding repeat containing 2(Xirp2) | 1.438311 | 9.54E-05 |
| Ankrd23 | ankyrin repeat domain 23(Ankrd23) | 1.394826 | 8.26E-07 |
| Scml4 | Scm polycomb group protein like 4(Scml4) | 1.358973 | 0.00146 |
| Hmgcs2 | 3-hydroxy-3-methylglutaryl-Coenzyme A synthase 2(Hmgcs2) | 1.32501 | 0.004116 |
| Col8a1 | collagen, type VIII, alpha 1(Col8a1) | 1.264303 | 9.46E-05 |
| Myh7 | myosin, heavy polypeptide 7, cardiac muscle, beta(Myh7) | 1.194301 | 0.005146 |
| Gsta1 | glutathione S-transferase, alpha 1 (Ya)(Gsta1) | 1.121673 | 0.000723 |
| Gck | glucokinase(Gck) | 1.10383 | 0.003587 |
| Abra | actin-binding Rho activating protein(Abra) | 1.101257 | 0.002577 |
| Cubn | cubilin(Cubn) | 1.079504 | 0.004336 |

| Gene ID | Gene name | log2FC | Padj |
|---|---|---|---|
| Rpl30-ps9 | ribosomal protein L30, pseudogene 9(Rpl30-ps9) | -5.45696 | 1.63E-05 |
| Lrrc4b | leucine rich repeat containing 4B(Lrrc4b) | -1.90947 | 4.39E-12 |
| Col6a5 | collagen, type VI, alpha 5(Col6a5) | -1.88152 | 0.00285 |
| Cngb3 | cyclic nucleotide gated channel beta 3(Cngb3) | -1.69794 | 3.07E-05 |
| Slc1a7 | solute carrier family 1 (glutamate transporter), member 7(Slc1a7) | -1.62561 | 0.003982 |
| Pycard | PYD and CARD domain containing(Pycard) | -1.47395 | 0.000234 |
| Gnmt | glycine N-methyltransferase(Gnmt) | -1.46677 | 0.00041 |
| Actg-ps1 | actin, gamma, pseudogene 1(Actg-ps1) | -1.32815 | 0.00124 |
| Lamb3 | laminin, beta 3(Lamb3) | -1.23098 | 0.00011 |
| Rpl19-ps1 | ribosomal protein L19, pseudogene 1(Rpl19-ps1) | -1.21323 | 0.000636 |
| Hsbp1l1 | heat shock factor binding protein 1-like 1(Hsbp1l1) | -1.1962 | 0.000583 |
| Plxnb1 | plexin B1(Plxnb1) | -1.14478 | 5.51E-06 |
| Ramp1 | receptor (calcitonin) activity modifying protein 1(Ramp1) | -1.11027 | 0.002142 |
| Rgs2 | regulator of G-protein signaling 2(Rgs2) | -1.07221 | 0.000164 |
| Epop | elongin BC and polycomb repressive complex 2 associated protein(Epop) | -1.05284 | 0.000409 |
| Tfpi | tissue factor pathway inhibitor(Tfpi) | -0.98374 | 3.39E-06 |
| Slc7a1 | solute carrier family 7 (cationic amino acid transporter, y+ system), member 1(Slc7a1) | -0.98253 | 8.98E-08 |
| Ciart | circadian associated repressor of transcription(Ciart) | -0.9362 | 3.03E-05 |
| Tmem100 | transmembrane protein 100(Tmem100) | -0.9299 | 0.000419 |
| Pfkfb1 | 6-phosphofructo-2-kinase/fructose-2,6-biphosphatase 1(Pfkfb1) | -0.92513 | 0.002135 |
| Mylk4 | myosin light chain kinase family, member 4(Mylk4) | -0.90474 | 0.003237 |
| Cep128 | centrosomal protein 128(Cep128) | -0.8961 | 0.00041 |
| Ccl11 | C-C motif chemokine ligand 11(Ccl11) | -0.86089 | 0.011554 |
| Kyat3 | kynurenine aminotransferase 3(Kyat3) | -0.85559 | 3.18E-09 |

**Supplemental Figure 5. Metabolomic profile of HFpEF hearts.** Targeted metabolomics from heart tissue focused on A) organic acids, B.) TCA cycle intermediates and C) medium and long chain acylcarnitine species (n=5-7). Data displayed as mean ± SEM. Unpaired, 2-tailed Student's t-test, with p values as displayed or * <0.05.

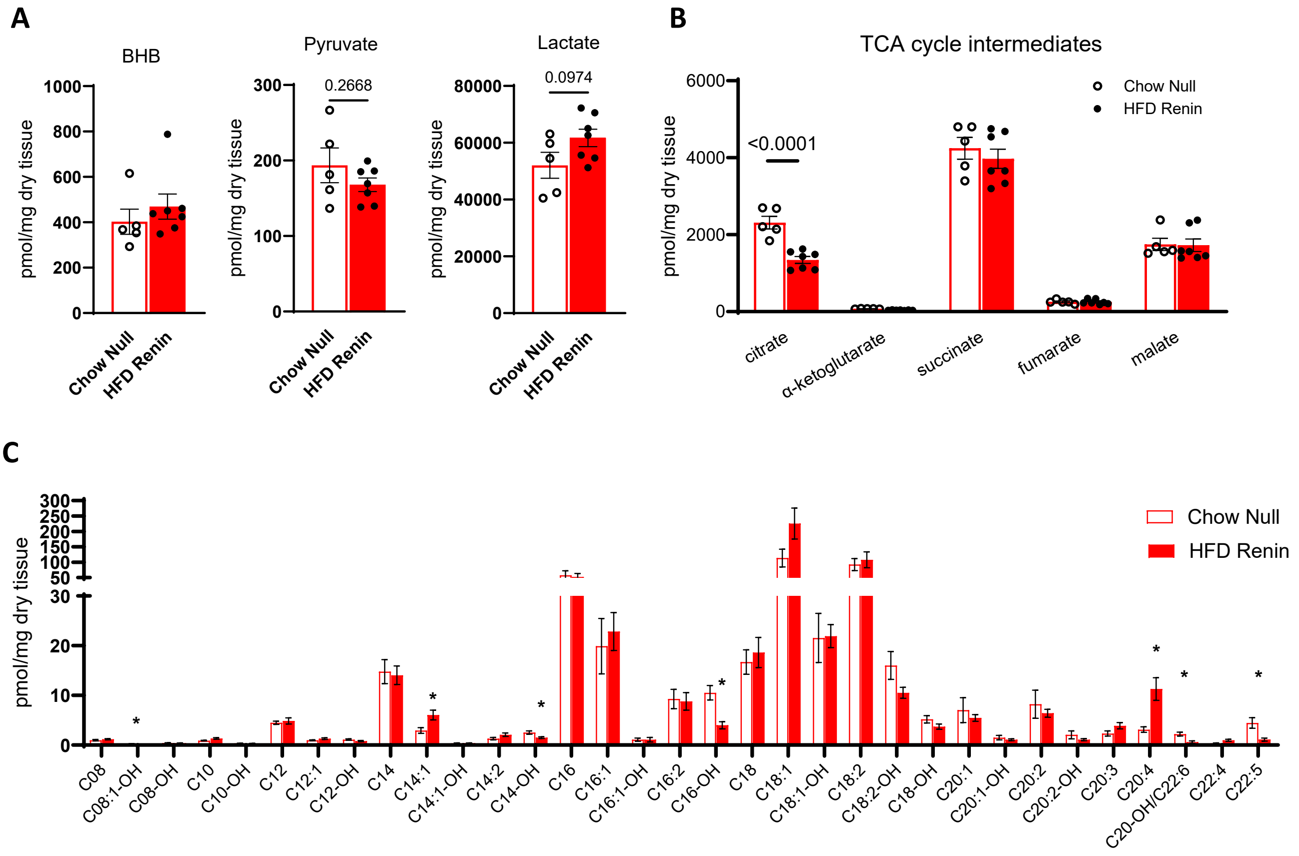

**Supplemental Figure 6. Effect of SGLT2i on mouse HFpEF (Penn).** A.) Normalized body weight of male control, HFpEF, and empa-treated HFpEF mice (n=15-27). B.) Echocardiographic measures of systolic function and strain. C.) LV measurements including echo-based LV mass indexed to body weight (LVMI) and tibia length normalized biventricular weight (BiV/TL). D.) Diastolic parameters including E/e', isovolumic relaxation time (IVRT) and left atrial (LA) enlargement. E.) Maximal exercise testing. Data displayed as mean ± SEM. One-way analysis of variance (ANOVA) followed by multiple-comparisons test. P values < 0.05 displayed on graphs.

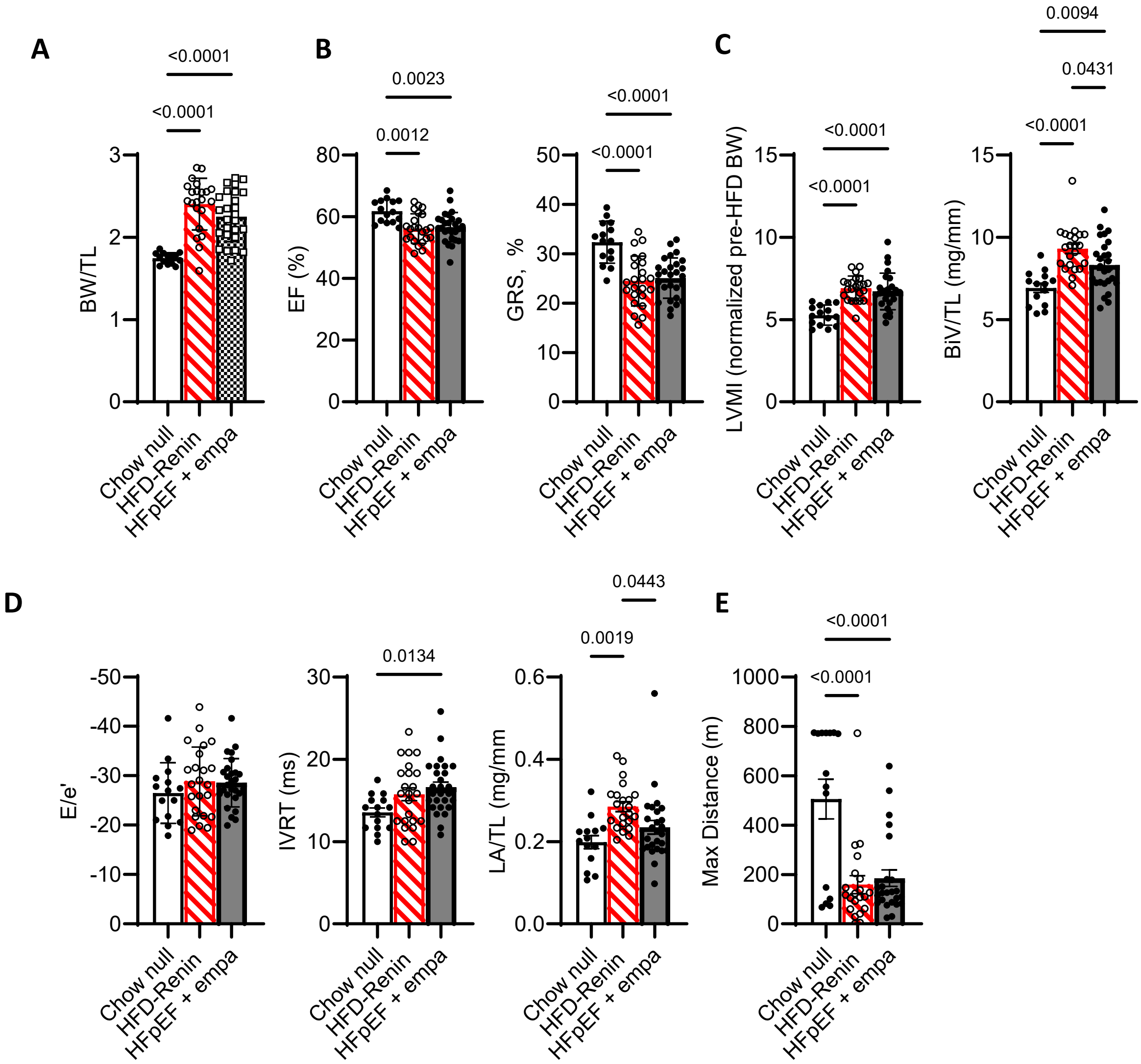

**Supplemental Table 1. Primer sequences.**

| | Forward | Reverse |
|---|---|---|
| *Acta1* | GTCCCAGACATCAGGGAGTAA | TCGGATACTTCAGCGTCAGGA |
| *Nppa* | GCTGCTTTGGGCACAAGATAG | GCAGCCAGGAGGTCTTCCTA |
| *Nppb* | GCTGCTTTGGGCACAAGATAG | GCAGCCAGGAGGTCTTCCTA |
| *Col1* | AGTCGCTTCACCTACAGCAC | TTCGATGACTGTCTTGCCCC |
| *Col3* | ACGTAAGCACTGGTGGACAG | GGAGGGCCATAGCTGAACTG |
| *Ccn2* | AGACCTGTGCCTGCCATTAC | ACGCCATGTCTCCGTACATC |
| *Elastin* | CTGGATCGCTGGCTGCATCCA | GTCCAAAGCCAGGTCTTGCTG |
| *Lox* | AAGCAGAGCCTTCCTGCAAA | GGTCACAGCGGTCTCGTTGT |
| *Postn* | TGGTATCAAGGTGCTATCTGCG | AATGCCCAGCGTGCCATAA |
| *Tgfb1* | CACCTGCAAGACCATCGACA | CATAGTAGTCCGCTTCGGGC |
| *Tgfb2* | CTTCGACGTGACAGACGCT | GCAGGGGCAGTGTAAACTTATT |
| In-Fusion and Kozak insertion | CCCTATATATGGATCgccaccatggacagaaggaggatgcc | CATTGCAATTGGATCttagcgggccaaggcgaat |
| Mutagenesis primer | AAGAACAAAGAGGAGTTCCTTGACCAATCTTACCTCCCC | CTCCTCTTTGTTCTTACGCCCCATTCAGCACTGAG |
| Sequencing primers | TGTCTACTACAACAGGGGTTCC | TTGCCTTGATAATGCTGCGG |

**Supplemental Table 2. Echocardiographic and gross morphologic parameters for HFD+Renin HFpEF model.**

| | 6 weeks – post AAV | | | |
|---|---|---|---|---|
| Parameter | Chow Null | Chow Renin | HFD Null | HFD Renin |
| Body weight (g) | 27.73±1.8 | 28.67±2.5 | 43.01±5.8*# | 38.21±4.7*# |
| Tibia length (mm) | 18.1±0.2 | 18.3±0.3 | 18.7±0.9 | 18.1±0.4 |
| BW/TL | 1.55±0.09 | 1.57±0.11 | 2.29±0.31* | 1.97±0.25*# |
| BiV weight (g) | 111.6±11.5 | 136.9±17.3 | 120.1±11.8 | 157.8±43.3 |
| BiV/TL | 6.17±0.91 | 7.48±0.86 | 6.41±0.57 | 8.70±2.2 |
| Wet Lung (mg) | 170.8±14 | 177±17 | 182±13 | 180±20 |
| Wet lung/TL | 9.44±0.79 | 9.69±0.92 | 9.71±0.67 | 9.92±1.1 |
| Wet/Dry Lung | 4.34±0.31 | 4.21±0.26 | 3.87±0.29 | 3.95±0.2* |
| HR (bpm) - M-mode | 578±45 | 570±35 | 612±67 | 552±31 |
| HR (bpm) - MVI/TDI | 467±52 | 465±16 | 487±28 | 433±51 |
| LVIDs | 2.50 0.40 | 2.04 0.41 | 2.35 0.29 | 2.25 0.47 |
| LVIDd | 3.79 0.35 | 3.40 0.34 | 3.60 0.19 | 3.47 0.42 |
| EDV | 49.6 8.9 | 36.3 8.9* | 40.9 6.7 | 42.8 11.6 |
| ESV | 19.6 6.2 | 12.5 5.2* | 19.2 2.6ǂ | 18.3 5.4ǂ |
| IVSd (mm) | 0.91 0.05 | 1.17 0.12* | 1.02 0.11 | 1.20 0.05*# |
| LVPWd (mm) | 0.82 0.04 | 1.06 0.16* | 0.94 0.09 | 1.09 0.14*# |
| RWT | 0.459 0.06 | 0.666 0.14* | 0.549 0.03 | 0.669 0.09*# |
| LVM | 117.5 12.78 | 143.7 11.56 | 131.5 17.66 | 158.9 45.8* |
| LVMI (by pre-diet BW) | 5.31 .076 | 5.85 0.44 | 5.46 0.90 | 6.97 1.86*# |
| FS | 34.5 5.8 | 40.6 7.7 | 34.9 5.8 | 35.6 8.3 |
| EF | 60.6 6.8 | 66.2 7.5 | 52.5 3.3ǂ | 55.7 5.6ǂ |
| E/A | 1.19±0.12 | 1.17±0.08 | 1.08±0.16 | 1.19±0.16 |
| E/e' | -22.66±2.96 | -28.41±6.86 | -26.4±6.21 | -31.15±6.0* |
| Tei Index | 0.42±0.08 | 0.47±0.1 | 0.48±0.1 | 0.47±0.07 |
| GLS | -17.8±2.2 | -15.3±3.7 | -15.0±3.1 | -13.4±3.6* |
| SR, long sys | 6.77±1.2 | 6.63±2.3 | 5.95±1.2 | 5.56±1.4 |
| SR, long dias | 8.03±2.2 | 8.46±2.7 | 6.67±1.1 | 5.97±1.9ǂ |
| GRS | 32.84±5.6 | 32.47±5.7 | 26.08±3.5*ǂ | 26.32±5.2*ǂ |
| SR, rad sys | 9.58±1.8 | 10.2±1.7 | 8.10±1.1ǂ | 8.27±1.8ǂ |
| SR, rad dias | 10.7±2.1 | 11.7±2.0 | 8.61±1.7ǂ | 8.01±2.6*ǂ |
| LA weight (mg) | 3.039±0.40 | 3.819±1.05 | 3.695±1.19 | 4.719±1.52* |
| LA/TL | 0.168±0.02 | 0.208±0.06 | 0.198±0.07 | 0.260±0.08* |
| LA min vol | 6.34±1.25 | 7.13±1.35 | 6.18±1.70 | 8.14±2.67 |
| LA max vol | 11.78±1.5 | 11.08±1.8 | 9.93±2.6 | 12.36±3.4 |
| LA strain | 22.7±11.8 | 13.5±5.8 | 18.2±6.8 | 11.2±5.0* |
| LA SRs | 5.41±2.3 | 4.13±1.4 | 5.99±2.3 | 3.12±1.0# |
| LA SRe | 7.35±4.0 | 5.13±2.1 | 7.34±2.3 | 4.14±1.0# |
| PV, S pk | 252.4±34.9 | 222.2±42.9 | 227.4±41.9 | 177.9±18.7ǂ |
| PV S mn | 149.8±18.5 | 132±31.5 | 130.8±21.1 | 103.7±8.7 |
| PV, D pk | 647.2±52.0 | 671.6±76.2 | 622.5±62.6 | 582±59.3ǂ |
| PV, D mn | 394.6±14.2 | 390.4±44.8 | 373±44.0 | 346±57.0 |

Values are means ± SEM; n = ***. Comparisons made by two-tailed Student's t test (2 conditions) or one-way ANOVA with Bonferroni post hoc correction (4 conditions). Statistical sig: * vs Chow null, # vs HFD null, ǂ vs chow renin

Abbreviations: BW, body weight; BiV, biventricular; BPM, beats per minute; EDV, end diastolic volume; ESV, end systolic volume; EF, ejection fraction; FS, fractional shortening; GLS, global longitudinal strain; GRS, global radial strain; HR, heart rate; IVSd, intraventricular diastolic septal thickness; LA, left atria; LVIDd/s, left ventricular internal diameter, end diastole/end systole; LVM, left ventricular mass; LVMI, indexed left ventricular mass; LVPWd, left ventricular posterior wall diastolic dimension; MVI/TDI, mitral valve inflow/tissue Doppler index; PV, pulmonary vein; RWT, relative wall thickness; SR, strain rate; TL, tibia length.
